# N-formylkynurenine but not kynurenine enters a nucleophile-scavenging branch of the immune-regulatory kynurenine pathway

**DOI:** 10.1101/2024.11.15.623703

**Authors:** Yongxin Wang, Euphemia Leung, Petr Tomek

## Abstract

Tryptophan catabolism along the kynurenine pathway (KP) mediates key physiological functions ranging from immune tolerance to lens UV protection, but the contributory roles and chemical fates of individual KP metabolites are incompletely understood. This particularly concerns the first KP metabolite, N-formylkynurenine (NFK), canonically viewed as a transient precursor to the downstream kynurenine (KYN). Here, we challenge that canon and show that hydrolytic enzymes act as a rheostat switching the fate of NFK between the canonical KP and a novel non-enzymatic branch of tryptophan catabolism.

In the physiological environment (37°C, pH 7.4), NFK deaminated into electrophilic NFK- carboxyketoalkene (NFK-CKA), which rapidly (< 2 minutes) formed adducts with nucleophiles such as cysteine and glutathione, the key intracellular antioxidants. Serum hydrolases suppressed NFK deamination as they hydrolysed NFK to KYN ∼3 times faster than NFK deaminates. Whilst KYN did not deaminate, its deaminated product (KYN-CKA) rapidly reacted with cysteine but not glutathione.

The new NFK transformations of yet to be confirmed functions highlight significance of NFK beyond hydrolysis to KYN and suggests the dominance of its chemical transformations over those of KYN in physiological environments. Enzyme compartmentalisation and abundance offer insights into the regulation of non-enzymatic NFK and KYN transformations that are emerging as contributors to immune regulation, protein modification, lens aging or neuropathology.

## 1. Introduction

Tryptophan is an essential amino acid acquired from the diet used for protein synthesis and bioactive metabolite production across 3 pathways (Fig. 1). In serotonin pathway, tryptophan is converted to serotonin and melatonin for regulation of a range of physiological and circadian rhythm functions^1-3^. Tryptophan can also enter the indole-pyruvate pathway, to produce indole metabolites that contribute to intestinal homeostasis and immunity^4, 5^. However, most of the dietary tryptophan (∼ 90%) enters the kynurenine pathway (KP) in the liver to maintain tryptophan homeostasis^6^. Two heam-containing dioxygenases, tryptophan 2,3-dioxygenase (TDO) and indoleamine 2,3-dioxygenase 1 (IDO1) are the principal catalysts of the first and rate-limiting step of the KP, the oxidation of tryptophan to N-formylkynurenine (NFK)^7-9^. Then, NFK is hydrolysed into kynurenine (KYN) by a hydrolase called arylformamidase^8, 10^. KYN can be further metabolised into neuro-modulatory metabolites such as anthranilic acid and kynurenic acid^11, 12^, and into the final product of KP, NAD^+^, a co-enzyme essential for redox reactions^13, 14^.

**Fig. 1.**
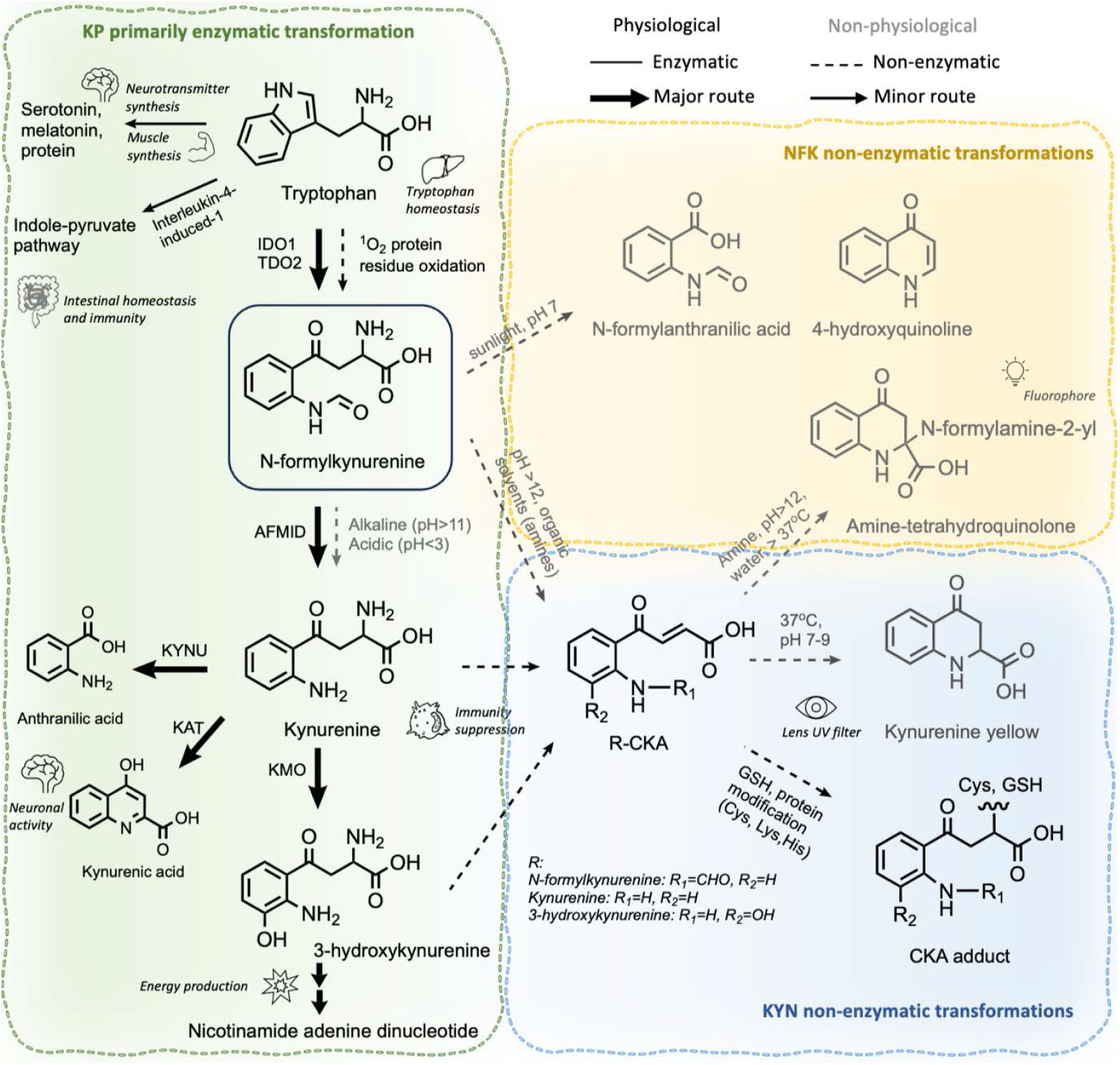
Enzymatic (solid line) and non-enzymatic (dashed line) transformations of NFK and KYN. Thick arrows represent major routes of transformation. Labels besides the arrows indicate assumed catalysts, if known. Metabolites and transformations associated primarily with physiological and non-physiological environments are black and grey, respectively. Italic labels of cartoon icons denote metabolites’ functions. Abbreviations: IDO1, indoleamine 2,3-dioxygenase 1; TDO2, tryptophan 2,3-dioxygenase 2; AFMID, arylformamidase; KYNU, kynureninase; KAT, kynurenine aminotransferase; KMO, kynurenine 3-monooxygenase; CKA, carboxyketoalkene; Cys, cysteine; Lys, lysine; His, histidine.

One of the key functions of the KP is arguably the regulation of immunity. KP contributes to maintaining immune tolerance during inflammation and infection^15-18^, pregnancy^19, 20^ and in immune-privileged sites such as cornea^21, 22^ and testes^23, 24^. Pro-inflammatory stimuli such as prostaglandin E2, tumour necrosis factor alpha, interferon gamma or lipopolysaccharide induce IDO1 expression^25, 26^ and extrahepatic KP, hence KP is typically inactive in most non-hepatic tissues. KYN appears to be the key culprit mediating KP’s immunosuppression and neutralising immunity primarily by activation of a transcription factor called aryl hydrocarbon receptor^27, 28^. It is now well established that most cancer types upregulate IDO1/TDO2 to accelerate KYN production and escape immune surveillance, that both worsen cancer patient prognosis and undermine immune checkpoint therapies^29-33^. Therefore, inhibiting KP enzymes by drugs has become a promising therapeutic avenue for potentiating cancer immunotherapy^34, 35^.

However, recent studies have started to unravel that non-enzymatic transformations of KP metabolites can also regulate physiological processes and immunity. Carreno et. al^36^ showed for the first time that elevated levels of a non-enzymatic deamination product of KYN, KYN- carboxyketoalkene (KYN-CKA), reduces inflammation pathology in mice with sickle cell disease. KYN-CKA has been known to react with nucleophilic protein residues (histidine, cysteine and lysine) of lens proteins and cause eye ageing^37, 38^. There is likely many more yet unidentified bioactive KP metabolites produced non-enzymatically, that can arguably limit the therapeutic potential of drugs targeting KP enzymes.

One potential source of these metabolites is NFK, the enigmatic first KP product that has been neglected for decades, omitted in tryptophan pathway diagrams, and viewed primarily as the precursor for enzymatic hydrolysis to KYN. But NFK seems to be more than meets the eye. The formyl group bestows NFK unique reactivity primarily through its transfer to diverse target molecules. At strongly alkaline pH > 11 in organic solvents, NFK deaminates into NFK-CKA, and its formyl group translocates to amines to form fluorophores^39, 40^. Similarly, the transfer of NFK’s formyl group to environmontal toxins was proposed to underpin enzyme-mediated detoxification in the liver^41, 42^. The NFK’s formyl group likely participates in one-carbon metabolism^43, 44^ and NFK itself has been suggested to activate immune-regulatory aryl hydrocarbon receptor but only at supraphysiological concentrations exceeding 1 mM^45^. Whilst these observations demonstrate potential physiological role of NFK, whether most NFK transformations play role *in vivo* remain unclear.

Here we report the identification of novel conjugates of NFK with biological nucleophiles that form at physiological conditions and represent a new non-enzymatic branch of tryptophan catabolism. The abundance of serum hydrolases regulates the flow between the canonical KP and the non-enzymatic NFK pathway suggesting biological importance of these non-enzymatic tryptophan by-products. Further, we highlight two important implications of these non-enzymatic transformation and sample processing for KP metabolomics.

## 2. Results

### 2.1 NFK stability in physiological and non-physiological environments

NFK’s main physiological fate is hydrolysis to KYN, which is accelerated either enzymatically^46^ or at pH extremes^40^ (Fig. 1). But we have found that in αMEM cell culture medium spontaneously alkalinised (pH 8.5) by overnight incubation at atmospheric CO_2_ pressure, NFK rapidly degraded (∼17 µM/hour) but not into KYN, as determined by high-performance liquid chromatography with a photodiode-array detector (HPLC-DAD). Similarly, at physiological conditions (pH 7.4, 37°C, 5% CO_2_), the majority of incubated NFK (100 μM) disappeared within 3 days but only 34.2 ± 2.4% hydrolysed to KYN (Fig. 2A, solid line). KYN was stable at both mildly alkaline (8.5 - 9) and physiological (7.4) pHs (Fig. 2A, dashed line). The physiological condition appears to transform NFK into metabolites other than KYN through pathways not involving KYN.

**Fig. 2.**
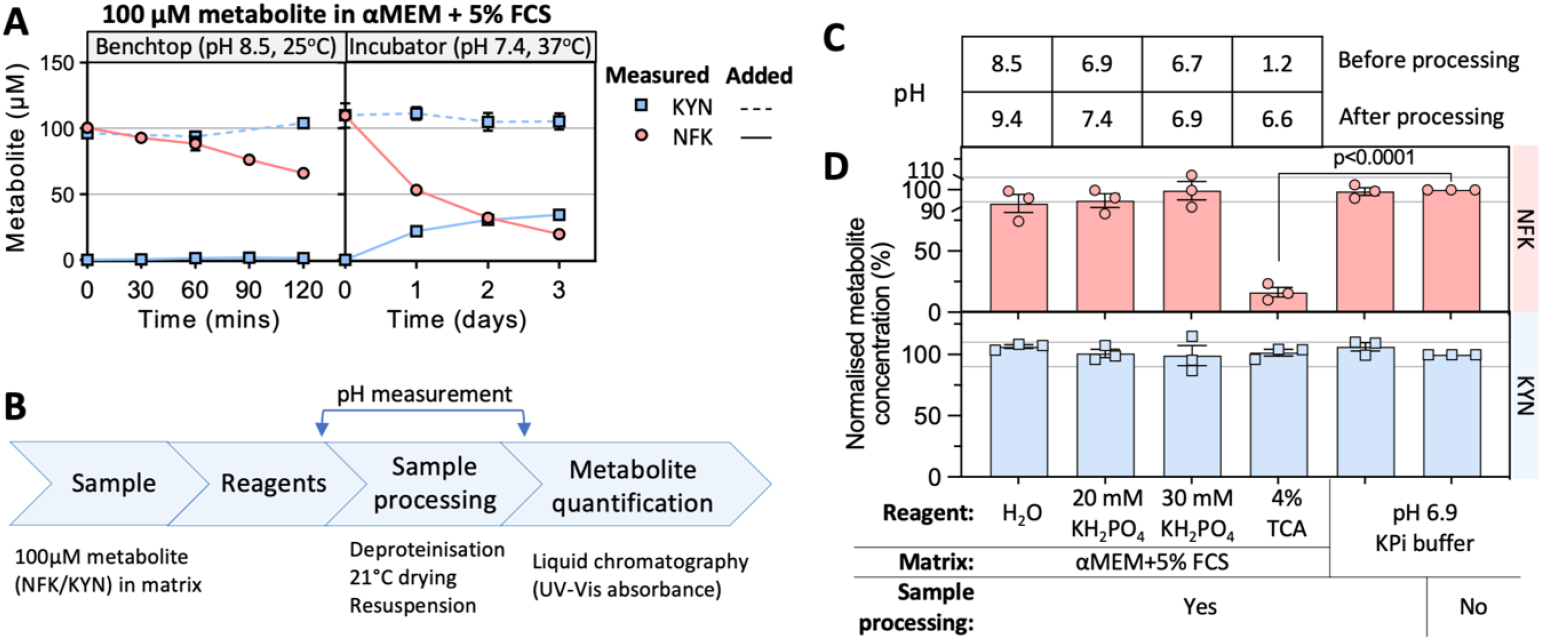
Mitigating non-enzymatic NFK transformations. **(A)** Stability of NFK (solid line) and KYN (dashed line) at conditions indicated above the graphs. Symbols and whiskers represent the mean and SEM of NFK (red circle) and KYN (blue rectangle) concentrations from 2 independent experiments. **(B)** The sample preparation procedure for generating data in Figs. 2C and 2D. **(C)** The mean sample pH before (top row) and after (bottom row) the sample processing (2 hours) from 3 independent experiments, measured by an electrode at 25°C, atmospheric CO_2_ pressure. The 4% TCA samples were neutralised by 375 mM K_2_HPO_4_ after the drying to aid solubility. **(D)** Concentration of NFK (pink) and KYN (blue) at different conditions (x-axis) normalised to that of metabolites stored on wet ice in pH 6.9 KPi buffer for the duration of sample processing. Bar heights and whiskers represent the mean and SEM from 3 independent experiments (symbols), respectively. Statistical analysis was conducted using Ordinary Two-way ANOVA, and the p-value was generated by Dunnett’s multiple comparisons test in GraphPad Prism software. The pH of αMEM+5% FCS matrix used to generate the results shown on this figure were not adjusted before preincubation. Abbreviations: αMEM: alpha Minimum Essential Medium; FCS: foetal calf serum; NFK: N-formylkynurenine; KYN: kynurenine; KPi: potassium phosphate buffer; TCA: trichloroacetic acid; SEM: standard error of the mean.

To accurately capture physiological NFK transformations without confounding degradation in the alkaline environment of sample processing, we have first tested whether adding non-volatile phosphate acid salt KH_2_PO_4_ could replace the volatile bicarbonate-CO_2_ buffering system of the cell culture medium, to stabilise neutral pH and mitigate metabolite degradation. We have used KH_2_PO_4_ concentrations (20 and 30 mM) comparable to that of the bicarbonate (∼ 26 mM) present in the αMEM culture medium, and monitored NFK and KYN stability and sample pH during ∼2-hour sample processing (Fig. 2B).

As hypothesised, KH_2_PO_4_ supplementation has completely overturned medium alkalinisation and promoted NFK stability outside of the cell culture incubator at atmospheric CO_2_ pressure. In contrast, 4% (w/v) trichloroacetic acid (a commonly used deproteinization agent for KP metabolomics)^47^ hydrolysed 83.7 ± 3.9% NFK to KYN (Fig. 2C and 2D). KYN was stable at all conditions tested (Fig. 2D). Sample processing at neutral pH 6.9 did not degrade NFK relative to wet-ice incubation (unprocessed sample), indicating that pH deviations from neutrality primarily drive NFK decomposition. The 30 mM KH_2_PO_4_ was deemed optimal to maintain neutral pH during sample processing due to its slightly better NFK stabilisation (∼10%) over the slightly weaker 20 mM KH_2_PO_4_ (Fig. 2C and 2D). Samples stabilised in KH_2_PO_4_ could be stored at -80°C for at least 38 days without significant NFK loss, thus allowing accumulation of samples for later analysis (Supplementary Fig. 1A). Having optimised the sample processing procedure, we then investigated the mysterious metabolic fates of NFK.

### 2.2 Catalysts of the NFK transformation

We posit that either physiological pH 7.4 alone and/or components of FCS or αMEM such as enzymes^48, 49^ and vitamins^50, 51^ catalyse NFK transformation. To identify these NFK metabolites and unravel their biosynthetic pathway, we incubated NFK (2 days, 37°C) separately in three components of the cell culture medium (H_2_O, αMEM, FCS) all adjusted to pH 7.4 by bicarbonate buffer, and analysed by HPLC-DAD and liquid chromatography - low-resolution mass spectrometry (HPLC-ESI-MS) (Fig. 3).

**Fig. 3.**
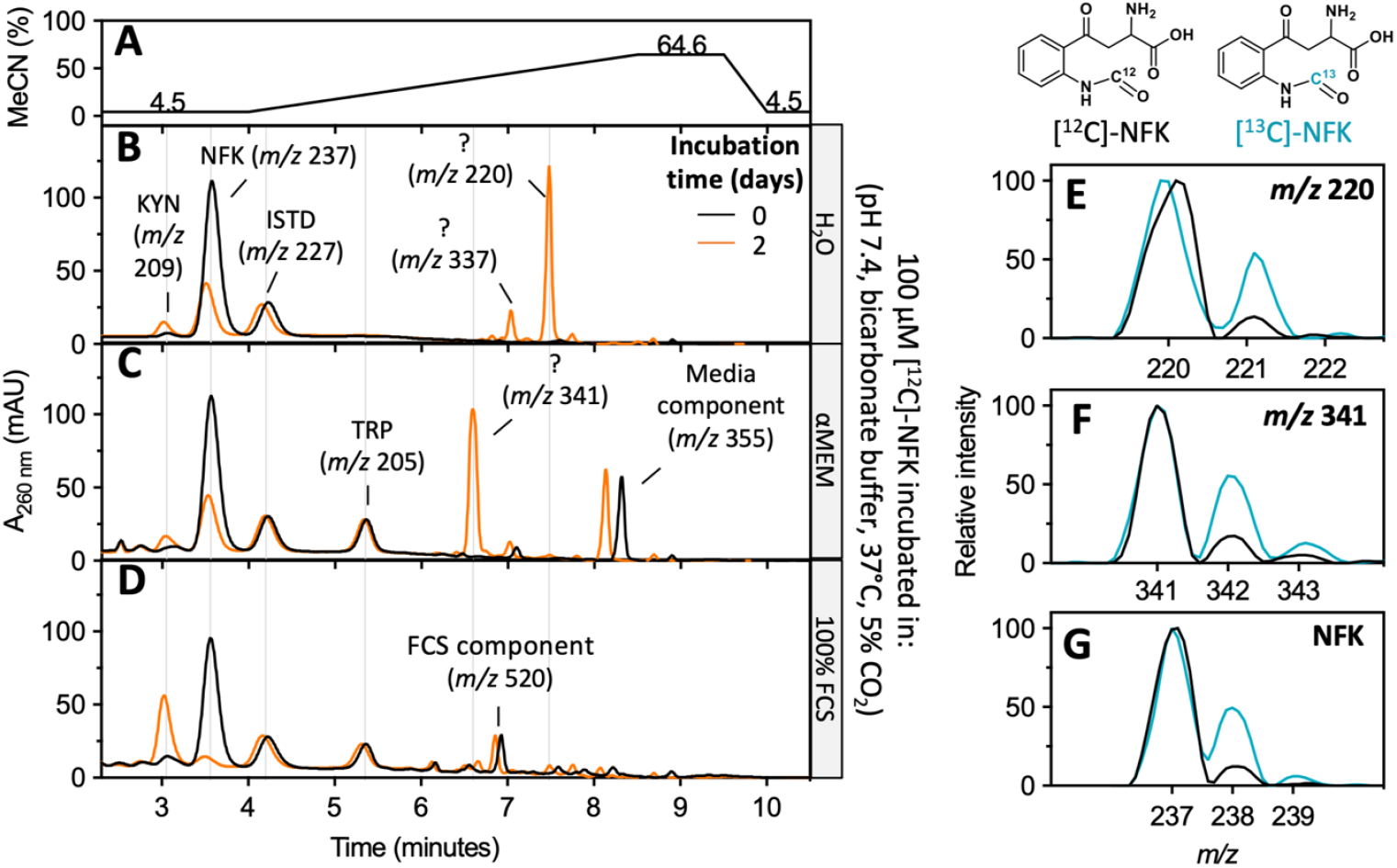
Unique NFK transformations driven by distinct components of cell culture medium. **(A)** Chromatographic gradient for separations in B-D. The time was adjusted for the HPLC dwell volume. The numerical labels above the gradient line indicate % MeCN. **(B-D)** HPLC-DAD (260 nm) chromatograms of NFK (100 µM) incubated for 0 (black) or 2 (orange) days in three matrices (rows; right-hand side annotations). Principal peaks (>10% of the NFK peak height) were labelled with their identity or ? if unknown, and mass-to-charge ratio (*m/z*) of the dominant ion. Absorbance and mass spectra for all annotated metabolites are in Supplementary Figs. 2 and 3, respectively. **(E-G)** Normalised mass isotope distribution of compounds *m/z* 220, 341 and NFK produced by incubation of ^12^C-NFK (black) or ^13^C-NFK (blue) in H_2_O (E,G) or αMEM (F). Abbreviations: ISTD: internal standard (3-nitro-L-tyrosine); TRP: tryptophan; MeCN: Acetonitrile.

Each of the 3 matrices produced a distinct dominant peak other than NFK, indicative of 3 unique and mutually exclusive NFK transformations (Fig. 3B - D). H_2_O produced a smaller and significantly more retained compound (*m/z* 220 [M+H]^+^, retention time 7.48 min), whereas αMEM generated a larger but slightly less retained entity (m/z 341 [M+H]^+^, retention time 6.6 min) in the absence of *m/z* 220 (Fig. 3B and 3C). Both *m/z* 220 and 341 were essentially absent in the FCS reaction where nearly all NFK was hydrolysed to KYN (Fig. 3D). Repeating the experiments with [^13^C-formyl]-NFK confirmed that *m/z* 220 and 341 carry the NFK’s formyl group (comparable ^13^C:^12^C isotope ratio), hence both are likely derived from NFK and not from KYN or other possible contaminants (Fig. 3E, 3F and 3G).

As the molecular weight difference between NFK (236 Da) and *m/z* 220 (219 Da) is consistent with that of ammonia (17 Da, NH_3_), we posit that *m/z* 220 is deaminated NFK, ie., NFK-CKA (Fig. 3B). Being a reactive electrophile, we further reasoned that NFK-CKA conjugates with a nucleophile present in αMEM to form *m/z* 341. Strikingly, theoretical conjugation of NFK- CKA with the only obvious nucleophile present in αMEM (cat. no. 12000063, ThermoFisher Scientific, Grand Island, NY, USA), amino acid cysteine (Cys; 568 μM), produces a conjugate of 340 Da expected to be the *m/z* 341 [M+H]^+^.

### 2.3 NFK deamination and addition to nucleophiles

To confirm that *m/z* 220 and 341 are NFK-CKA and NFK-Cysteine adduct, respectively, we have reacted purified intermediates (37°C, 5% CO_2_, 2 days) in bicarbonate-buffered H_2_O (pH 7.4) and analysed the reactions by HPLC-ESI-MS. The results have confirmed both NFK deamination and thiol conjugation. The authentic standard of NFK-CKA produced identical *m/z*, retention time, and UV spectrum as *m/z* 220 generated by incubation of NFK in pH 7.4 H_2_O (Fig. 4A and 4B). Similarly, *m/z* 341 formed in the reaction of purified NFK or NFK- CKA with 568 µM Cys (Figs. 4A - C). Reduced glutathione (GSH) also formed a NFK-adduct when reacted with NFK or NFK-CKA (Fig. 4B and 4C). This has confirmed NFK-CKA as the intermediate to NFK-GSH/Cys adducts and generalised NFK’s conjugation potential to biological thiols. Unexpectedly, at the same conditions that deaminated NFK into NFK-CKA, KYN remained stable (Fig. 4D) but KYN-CKA formed adducts with both Cys and GSH (Fig. 4E). This observation challenges the seemingly well-established reactivity of KYN with biological nucleophiles.

**Fig. 4.**
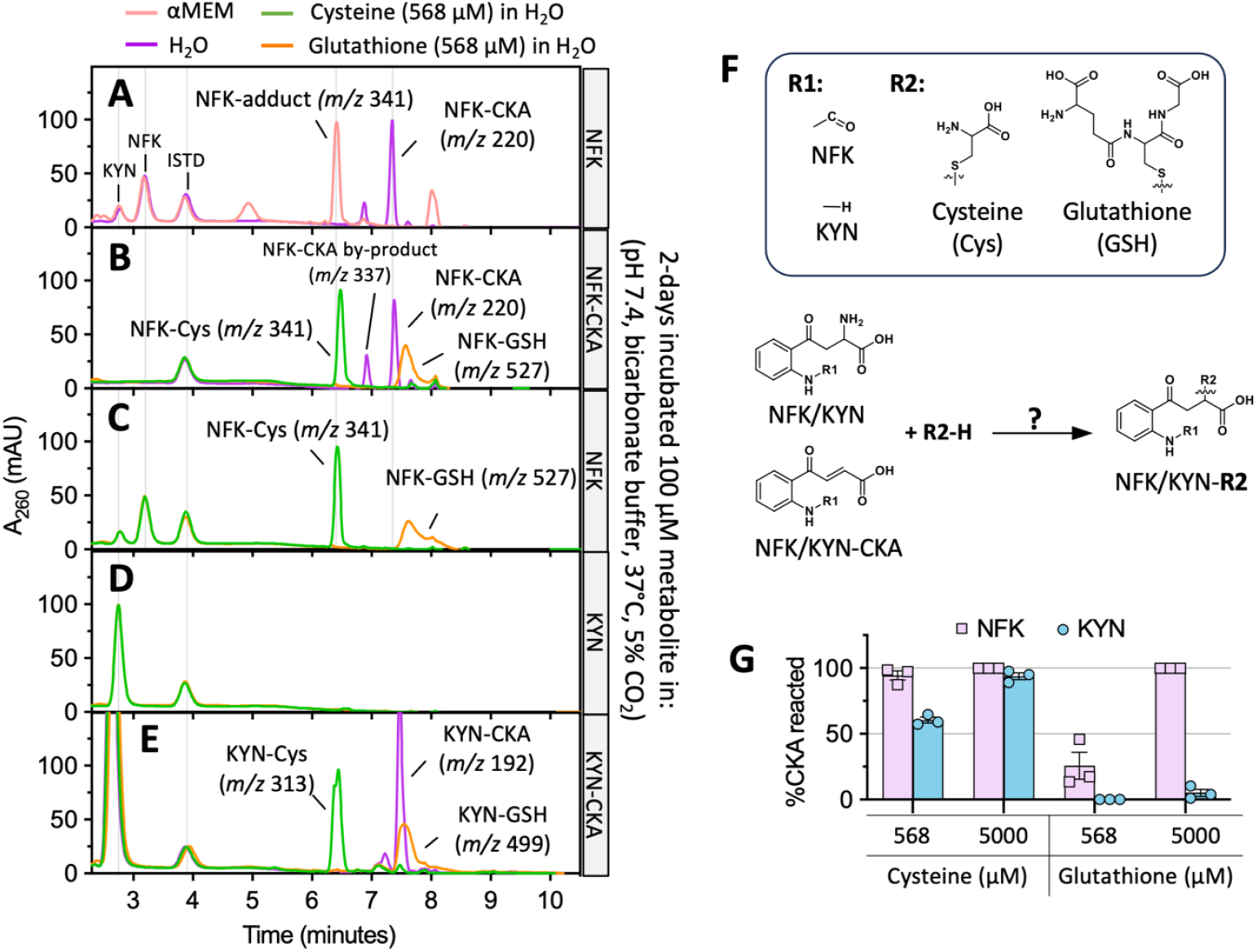
NFK but not KYN deaminates and conjugates to biological thiols. **(A-E)** HPLC-DAD chromatograms of each metabolite (rows), incubated for 2 days at 37°C with four different vehicle solutions or thiols (colours). Chromatograms were retention time-aligned by the internal standard (ISTD, retention time ∼3.9 mins). Peaks of interest were labelled with their identity and dominant *m/z* in their mass spectra. The absorbance and mass spectra of the labelled peaks are in Appendix Figs. 2 and 3, respectively. **(F)** Reactions tested in A-E **(G)** % loss of KYN (blue circle) and NFK (purple rectangle) CKAs relative to the vehicle control after a 2-min reaction with biological thiols in pre-incubated bicarbonate-buffered H_2_O (pH 7.4, ∼37°C). Bar heights and whiskers are mean and SEM of 3 independent experiments (symbols).

To test this notion, we have quantified reactivity of NFK-CKA and KYN-CKA with GSH and Cys in a 2-min reaction (37°C, pH 7.4) at concentrations mimicking median intracellular GSH (5 mM)^52, 53^ and αMEM Cys (568 µM) levels. NFK-CKA indeed reacted with thiols substantially faster than KYN-CKA (Fig. 4G). Cys was surprisingly more reactive than GSH and both CKAs reacted with it to completion except for KYN-CKA at low cysteine concentration. That was likely due to cysteine’s lower pKa (8.3 at 25°C)^54^ favouring thiol ionisation at physiological pH 7.4. GSH barely reacted with KYN-CKA at neither concentration but consumed all NFK-CKA within 2 minutes at 5 mM. This almost instantaneous consumption of NFK-CKA by GSH at the median intracellular concentration argues for the dominance of NFK deamination and concomitant reactivity *in vivo*. This would, however, prevent NFK from hydrolysing into KYN via the canonical KP. Therefore, a mechanism for diverting the flow between NFK hydrolysis and deamination must exist.

### 2.4 Regulation of NFK transformations

As FCS almost completely suppressed NFK deamination and thiol adduct formation in favour of NFK hydrolysis (see Fig. 3D), we reasoned that the abundance of FCS components acts as a rheostat regulating the hydrolysis and deamination of NFK. To test this concept, we have titrated FCS in αMEM and quantitated the rates of NFK (100 µM) transformations over the course of 3 days (pH 7.4, 37°C, 5% CO_2_). Increasing the FCS concentration indeed strongly accelerated NFK to KYN hydrolysis and at 100% FCS, the hydrolysis was more than 3 times faster than NFK deamination or hydrolysis in the absence of FCS (Fig. 5A and 5B). But at < 5% FCS, NFK deamination and thiol adduct formation rate trumped hydrolysis by the factor of 2 to 8 demonstrating that most NFK is not hydrolysed to KYN at low serum levels. The comparable rate of NFK deamination and Cys adduct formation further corroborated the rapid nature of the thiol conjugation (Fig. 5A - B).

**Fig. 5.**
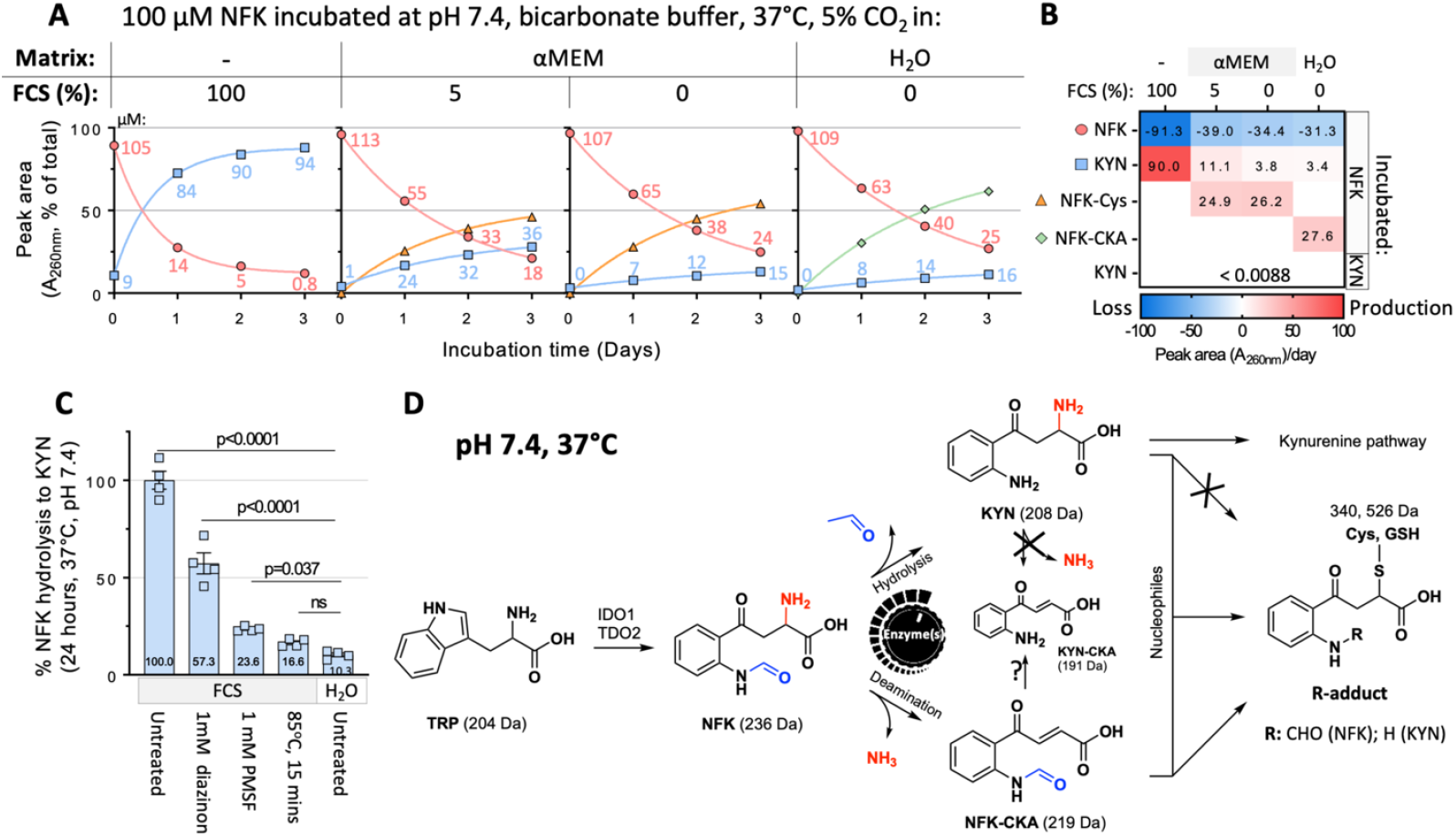
Serum hydrolases alter NFK’s fate between hydrolysis and deamination. **(A)** Time courses of NFK transformations 37°C in 4 different pH 7.4 bicarbonate-buffered solutions. Data points are internal standard-normalised peak areas expressed as % of metabolite sum at each incubation time. Points and whiskers are means and SEM of two independent experiments. Solid lines connecting the points are 1-phase decay non-linear regression fits. Numerical labels indicate μM NFK and KYN. **(B)** A heat map of initial reaction velocity (Peak area.day^-1^) of time courses in panel A (NFK) or Supplementary Fig. 1B (KYN) calculated between time 0 (t_0_) and reaction half-life (t_1/2_) as: (Peak area _t1/2_ – Peak area _t0_) / t_1/2_. **(C)** Sensitivity of NFK hydrolysis to pan-hydrolase inhibitors (PMSF and diazinon) and heat denaturation. Treatments (x-axis) were performed prior to adding NFK (100 μM). Bar heights and whiskers are mean and SEM, respectively, of 4 technical repeats (square points) across 2 independent experiments. Data points are KYN concentrations normalised to untreated FCS. The p-values were generated by Dunnett’s multiple comparisons test post 1-way ANOVA in GraphPad Prism. **(D)** Proposed integration and regulation of canonical KP and novel non-enzymatic NFK pathway. Question marks and crosses indicate hypothetical and unlikely reactions, respectively. Abbreviations: PMSF; phenylmethylsulfonyl fluoride.

Serum’s extraordinary efficiency in hydrolysing NFK to KYN prompted us to investigate whether the reaction is catalysed exclusively by enzymes. The near complete (83.4 ± 1.0%) inhibition of NFK hydrolysis by heat denaturation indicates that enzymes control NFK to KYN hydrolysis - at least in the bovine blood serum (Fig. 5C). Partial sensitivity (42.7 ± 5.4 % to 76.4 ± 0.6%) of NFK hydrolysis to pan-hydrolase/protease inhibitors diazinon and phenylmethylsulfonylfluoride further suggests those enzymes are hydrolases^10, 55-57^. We cannot exclude however, that NFK hydrolysis involves other classes of enzymes, as neither inhibitor reached the potency of heat denaturation (Fig. 5C). In sum, these data demonstrate that the abundance of hydrolytic enzymes determines NFK’s fate between NFK hydrolysis and deamination (Fig. 5D).

## 3. Discussion

We report a novel non-enzymatic branch of KP initiated by deamination of its first metabolite NFK into a highly reactive NFK-carboxyketoalkene that almost instantaneously forms conjugates with biological thiols at physiological levels and conditions (Fig. 4). The abundance of hydrolase enzymes appears to be the master regulator modulating NFK’s fate between the canonical hydrolysis along the KP and deamination along this novel nucleophile-scavenging pathway (Fig. 5). To our knowledge, this is the first report of NFK transformations other than its hydrolysis to KYN occurring at physiological conditions as NFK reactions have been observed primarily at non-physiological conditions^39, 40, 58^ such as extreme pH, supraphysiological concentrations or temperature (Fig. 1).

Whilst the function of this NFK branch remains unclear for now, one can argue it likely modulates redox equilibrium by scavenging key antioxidants such as glutathione. It is also possible that NFK-CKA conjugates nucleophilic protein residues such as cysteine, lysine or histidine. That is expected to alter the proteins’ function, and could contribute to eye aging as previously shown for other KP metabolites such as KYN and 3-hydroxy-KYN^37, 59-61^. In this situation, glutathione would act as a sacrificial nucleophile to neutralise NFK-CKA and protect lens crystallins from harmful modifications^62, 63^.

The most striking finding from our study was that, unlike NFK, KYN could not deaminate at physiological conditions (37°C, pH 7.4) even after 2 days of incubation at a supraphysiological concentration of 100 μM (Fig. 4D and 5B) which would be expected to substantially accelerate the reaction rate. This observation challenges the seemingly well-established KYN deamination into KYN-CKA *in vivo* and its subsequent conjugation to nucleophilic protein residues or free thiols^37, 38, 63, 64^. Our data suggest that NFK rather than KYN initiates and dominates non-enzymatic transformations in physiological environments, and that KYN derivatives form by subsequent enzyme-catalysed or spontaneous hydrolysis of NFK derivatives (Fig. 5D). NFK derivatives are thus expected to functionally overlap with other known CKAs *in vivo*, and play roles in post-translation modification^37, 61, 64-66^ and immunosuppression^36^.

Superior NFK deamination rate and NFK-CKA reactivity over that of other KP metabolites such as KYN, 3-hydroxykynurenine and 3-hydroxykynurenine glucoside^66^ support this notion. For example, in one study, only 20% of 3mM KYN deaminated in 4 days (pH 7.4, 37°C)^66^. In contrast, this work has demonstrated that at a 30-fold lower concentration (100 μM) and half the incubation time (2 days), substantially more NFK (46.3 ± 1.2%) deaminated into NFK- CKA (Fig. 3B and 5A). Similarly, median intracellular glutathione levels (5 mM)^52, 53^ consumed all NFK-CKA (100 μM) within 2 minutes at 37°C whereas only 7.2% KYN-CKA reacted at the same conditions. That demonstrates the marked impact of the formyl group on the molecule’s reactivity. Although Cys reactivity was more comparable between KYN and NFK CKAs, the tendency of KYN-CKA to react slower at lower Cys levels (Fig. 4G) suggests that at physiological cysteine concentrations (∼ 11 µM)^67, 68^, NFK-CKA will also likely surpass KYN-CKA. This would be highly relevant for modification of Cys residues that are relatively rare in mammalian proteins at less than 2.26 %^69, 70^. In sum, NFK-CKA appears to out compete KYN-CKA in scavenging both free antioxidants and likely even nucleophilic amino acid residues. But for NFK deamination to take place, canonical hydrolysis of NFK to KYN along KP must be less favourable. That is true as NFK deamination is ∼2 to 8 times faster than its hydrolysis but only when hydrolytic enzymes are scarce (Fig. 5A - B). Therefore, high hydrolase levels will invariably favour hydrolysis along the KP.

This would mean that hydrolytic enzyme abundance mediates a key switch between NFK deamination and hydrolysis along KP (Fig. 5D). If confirmed, this would represent an ingenuine endogenous way of compartmentalising non-enzymatic transformations as tissues and cells are expected to vary in hydrolytic enzyme content. For example, in extracellular fluids and blood consisting of up to 50% of hydrolysis-favouring serum^71, 72^, little NFK deamination is expected to occur and KP proceeds preferentially. Low NFK concentration (< 1 μM) in mammalian blood supports this theory^46, 73^. We cannot exclude though that low circulating NFK levels result from NFK not entering the bloodstream and being metabolized primarily in solid tissues. On the other hand, in tissues such as the eye’s lens, NFK is anticipated to preferentially enter the deamination rather than KYN pathway as the lens contains primarily structural non-catalytic proteins^74, 75^. But could this NFK deamination occur *in vivo*? We reason that is highly likely due to the presence of analogous CKAs of other KP metabolites such as KYN and 3-hydroxy-KYN in cultured cells, eye’s lens and other animal tissues^36, 38, 61, 76^. However, further experimentation will be necessary to address this question decisively.

Our findings also raise two important considerations regarding technical execution and interpretation of KP metabolomics studies. Firstly, TCA should be avoided during sample preparation. This strong organic acid has been a reagent of choice for decades for quantification of KYN in absorbance/fluorescence microplate assays^77-80^ and for sample deproteinization prior to bioanalytical measurements^81-83^. However, TCA rapidly hydrolyses NFK into KYN (NFK half-life ∼ 9 minutes at 37°C in 5% (w/v) TCA; data not shown). This alters the original KYN and NFK abundance in the sample, confounds interpretations and likely explains NFK’s absence in many KP studies^84, 85^. Where possible, we recommend processing both NFK and KYN at a neutral or slightly acidic pH between 6 to 7 where NFK appears most stable (Fig. 2). The pH-stabilising method developed in this study mitigates NFK degradation during sample processing from cell culture media and will be applicable to other carbonate-buffered systems such as plasma or animal tissues. This allows safe and reliable sample accumulation and processing in large batches without the need for deproteinisation beforehand (Fig. 2, Supplementary Fig. 1A). Secondly, spontaneous NFK transformations to KYN and other derivatives in cell culture medium have potential to misconstrue inferences about cellular KP obtained from secretome measurements^80, 86-88^. In another words, NFK and KYN concentrations in cell culture medium may not accurately reflect intracellular KP metabolism. For critical experiments, it is advisable to conduct intracellular KP metabolome measurements during the course of minutes to hours using stable isotope labelled tryptophan to overcome this limitation^89, 90^.

In conclusion, the identification of a novel branch of KP, NFK transformations and hydrolase- based regulation highlights there is still much to learn about the fates of tryptophan metabolites, particularly the non-enzymatic ones that have already started to be recognised as key actors in regulation of immunity^36^, lens aging^37, 61^, and neuropathology^91^. NFK is clearly not just a mere precursor to KYN as has been thought for decades. Its unique reactivity significantly exceeds that of its more famous cousin KYN even in physiological conditions. That makes NFK likely a key initiator of non-enzymatic KP transformations. The fascinating insight that hydrolytic enzymes alter the flow between enzymatic (KP) and non-enzymatic (thiol-scavenging) pathway (Fig. 5D) provides clues on how these non-enzymatic branches of metabolism can be regulated. Whilst comprehensive understanding of regulation and function of NFK thiol- scavenging pathway are yet to be demonstrated, we hope that our pH-stabilising method will facilitate and expedite novel chemical biology discoveries about this enigmatic tryptophan metabolite and other derivatives long overlooked in the literature.

## 4. Materials and methods

### 4.1 Chemicals and reagents

Milli-Q water (EQ7000 Ultrapure Water Purification System, Merck KGaA, Darmstadt, Germany) was used to prepare all aqueous solutions. NFK (98.2% purity by manufacturer, 96% purity by in-house HPLC-DAD) was purchased from PharmaCore CO., LIMITED (Kunshan, China). Formyl-^13^C NFK was synthesised as published previously^40^. Crude KYN-CKA (20% purity by HPLC-DAD) and NFK-CKA (5.2% purity by UV/VIS absorbance, ε /M^-1^ cm^-1^ 1584 at 322 nm) were prepared as described previously^40^. L-kynurenine (≥98%, cat. no. K8625), L-Cysteine hydrochloride (≥98%, cat. no. C7477), L-Glutathione (≥98%, cat. no. G4251), phenylmethanesulfonyl fluoride (≥98.5%, cat. no. P7626), K_2_HPO_4_ (≥98%, cat. no. P3786) and diazinon (cat. no. 45428) were all purchased from Sigma-Aldrich (St Louis, MO, USA). KH_2_PO4 (cat. no. 1.04873.1000) and NaOH (cat. no. 1.06498.0500) were from Merck KGaA (Darmstadt, Germany). Hydrochloric acid (cat. no. AC07742500) was from Scharlau (Barcelona, Spain). Foetal calf serum (cat. no. 38827105) was from Moregate Biotech (Hamilton, New Zealand). Alpha MEM cell culture medium (αMEM, cat. no. 12000063) was from Thermo Fisher Scientific (Grand Island, NY, USA). Trichloroacetic acid (cat. no. 04141- 500G) was from ECP Labchem (Auckland, New Zealand). Glacial acetic acid (cat. no. AJA1- 2.5L) was from Ajax FineChem (Auckland, New Zealand). All organic solvents were HPLC grade. Acetonitrile (>99.9%, cat. no. A955-4) and MeOH (> 99.9%, cat. no. A456-4) were from Fisher Chemical (Pittsburgh, PA, USA). NaHCO_3_ (cat. no. 102474V), sodium acetate trihydrate (cat. no. 10235), and ammonium formate (cat. no. 100246D) were from BDH Chemicals Ltd (Poole, England). Formic acid (>98%, cat. no. 1.00264.1000) was from Merck KGaA (Espoo, Finland). Potassium phosphate (KPi; 400 mM) buffer stock solution was prepared by mixing equal volumes of 400 mM K_2_HPO_4_ and KH_2_PO_4_ in Milli-Q water and adjusted to pH 6.9.

### 4.2 Experimental

#### 4.2.1 General experimental protocol and matrix preparation

Unless stated otherwise, all experimental incubations were performed in 96-well flat-bottom transparent polystyrene microplates (cat. no. 655101, Greiner, Frickenhausen, Germany) and processed for HPLC analysis (section 4.2.2) except for Fig. 5C reactions that were conducted in a polypropylene HPLC insert (cat. no. 4025P-631, JG Finneran; Vineland, NJ, USA) immediately prior to the HPLC acquisition (section 4.3.1). The general procedure consisted of i) incubating reaction matrices overnight (∼12 to 14 hours; 37°C, 5% CO_2_) with or without non-KP reagents (pan-hydrolase inhibitors PMSF or diazinon, and thiols GSH or Cys) to equilibrate temperature and pH, and ii) initiating the experiment by combining a matrix aliquot (110 uL) with KP metabolites (10 μL; final concentration 100 μM) as specified in each respective figure/legend.

The reaction matrices were: αMEM, αMEM+5% FCS, 100% FCS, and H_2_O supplemented with buffering components (26 mM NaHCO_3_ and 1 mM KH_2_PO_4_) present in αMEM. Except for experiments in Fig. 2, all matrices were adjusted by 0.4 M HCl to maintain pH 7.4 for the duration of the experiment. pH measurements were conducted by a glass micro electrode (cat. no. 8220BNWP, ThermoFisher Scientific, Australia) connected to the Orion Star™ A211 Benchtop pH Meter (ThermoFisher Scientific). After the pH adjustment, the final concentration of NaHCO_3_ in the three matrices was calculated to be 14.4 mM (αMEM, αMEM + 5% FCS) and 10.6 mM (H_2_O) based on molar equivalents of HCl added. In Fig. 5C experiments, heat-denatured FCS matrix was a supernatant obtained after spinning (15,000 *g*, 15 minutes, 4 °C) 100% FCS (200 uL; 1.7 mL Eppendorf tubes) heated at 85°C for 15 minutes in a Thermomixer Comfort heating block (Eppendorf, Hamburg, Germany). Pan-hydrolase inhibitors were added to the 100% FCS matrix from 200 mM neat DMSO stock solutions (final concentration 1 mM in 0.5% (v/v) DMSO).

#### 4.2.2 Sample preparation for HPLC analyses

At each designated time point, samples were aliquoted (80 μL) into a 96-well polystyrene microplate and, to stabilise pH between 6.5 to 6.9, 400 mM KH_2_PO_4_ was added to matrices to a final concentration of 15.4 mM (αMEM, αMEM+5% FCS, and H_2_O), 23 mM (100% FCS), or 20 or 30 mM (Fig. 2 experiments that used pH-unadjusted aMEM + 5% FCS).

Samples were stored at -80°C or analysed immediately. KYN and NFK calibration curves were prepared in KPi buffer (10 mM, pH 6.9) in duplicates of four 5-fold serially diluted concentrations from 125 uM to 1 uM to cover the range between the HPLC-DAD’s detection limit (∼1 µM) and the expected maximum metabolite concentration. Calibrations were prepared fresh for each HPLC acquisition batch and treated identically as the samples.

After thawing (25°C, 15 to 20 minutes, room temperature), if necessary, samples were aliquoted (50 μL) into a 96-well polypropylene V-bottom plate (cat. no. 651201, Greiner, Frickenhausen, Germany) or tubes (FCS-containing samples) containing 10 μL internal standard (3-nitro-L-tyrosine, 240 μM in 30 mM pH 6.9 KPi buffer). FCS-containing samples were then deproteinized by receiving 200 uL ice-cold methanol (80% v/v final concentration) and incubated on wet ice for 30 minutes, spun (15,000 *g*, 15 minutes, 4°C, Heraeus Multifuge 3S-R, Thermo Fisher Scientific, Germany), supernatants evaporated to complete dryness (21°C, 2 to 2.5 hours, 10 - 30 mbar) in a centrifugal vacuum concentrator (CentriVap, LABCONCO, Kansas City, MO, USA), and resuspended in 45 µL Milli-Q water. Samples (38 to 40 μL) were subsequently transferred to polypropylene HPLC inserts (cat. no. 4025P-631, JG Finneran; Vineland, NJ, USA). The remaining samples (5 to 7 μL for each individual sample) were pooled as a quality control (QC) sample. Inserts were arrayed in a 96-well deep well plate (cat. no. 780270, Greiner, Frickenhausen, Germany), spun (200 *g*, 1 minute, 4°C, Heraeus Multifuge 3S-R) to remove air pockets, and placed in amber glass HPLC vials.

### 4.3 Metabolite quantification and identification on bioanalytical instruments

Samples (35µL) were kept in a 5°C autosampler prior to analysis and injected onto a reverse-phase Zorbax SB-C18 column (150×3 mm, 5 μm particle size, Agilent, Santa Clara, CA, USA, cat. no. 883975-302) eluted by a binary mobile phase gradient at 0.65 mL/min flow rate and 30°C. Metabolites were analysed either by DAD or mass spectrometry (MS) detector as specified below. The QC sample was injected at regular intervals 3 to 4 times during each HPLC acquisition batch (8 – 12 hours) to assess metabolite stability in the 5°C autosampler and instrument performance. No marked degradation of any measured metabolite was observed on autosampler storage.

#### 4.3.1 Liquid chromatography - diode array detector (HPLC-DAD)

An Agilent 1100 Series HPLC Value System was used to acquire UV chromatograms. The mobile phases consisted of A = 80% acetonitrile and B = 20 mM sodium acetate (pH 4) supplemented with 3% acetonitrile. They were mixed as follows: 0 - 2 min (2% A), 6.5 – 7.5 min (80% A), 8 – 10.5 min (2% A). NFK and KYN were quantified from 320 nm and 360 nm chromatograms, respectively, as ISTD-normalised peak areas by comparing to the calibration curve.

#### 4.3.2 Liquid chromatography - low-resolution mass spectrometry (HPLC-ESI-MS)

To identify molecular masses of metabolites, an Agilent 1290 HPLC/6460 series liquid chromatography - triple quadrupole JetStream ESI mass spectrometry system was used. The mobile phases consisted of A = 0.1% (v/v) formic acid in Milli-Q (pH ∼ 2.8) and B = 80% (v/v) acetonitrile. They were mixed as follows: 0 - 2 min (94.3% A), 6.5 – 7.5 min (19.2% A), 8 – 10.5 min (94.3% A). The acquisition conditions were: 135-1000 m/z range, 500 scans/second, 100 V fragmentor voltage, 7 V cell accelerator voltage. Ionisation conditions were: 300°C drying gas temperature, 10 L/min gas flow. 45 psi nebulizer pressure, 3500 V positive/negative capillary voltage, positive/negative 500 V nozzle voltage.

### 4.4 Data analysis

Mass spectrum data were analysed with Agilent MassHunter software (Version 10.0, Agilent, Santa Clara, CA, USA). Microsoft software (Excel and PowerPoint, Version 16.0, Microsoft, Redmond, WA, USA) and GraphPad Prism (Version 9.3.1, Dotmatics, Boston, MA, USA) were used to analyse data and plot figures.

## Acknowledgements

We would like to thank Mr Sree Sreebhavan and Mr Sisira Kumara from Auckland Cancer Society Research Centre (University of Auckland, New Zealand) for technical assistance with bioanalytical instruments.

## Funding support

- School of Medical Sciences International Masters Scholarship (University of Auckland, Auckland, New Zealand) to YW.
- Royal Society of New Zealand’s Marsden Fast-start grant (MFP-UOA-2201) to PT.
- Auckland Medical Research Foundation’s Project Grant (1122012) to PT and EL.
- Cancer Research Trust New Zealand’s Project Grant (CRTNZ 2168 RPG) to PT.
- Core technical and salary support from Auckland Cancer Society Research Centre, Cancer Society Auckland Northland and School of Medical Sciences.

## Author’s contribution

YW designed, conducted, analysed and interpreted the experiments, and drafted the manuscript. EL contributed to experimental design, supervision and data analysis. PT conceived, designed, directed and supervised the study, and contributed to experimental design and execution. All authors contributed to the manuscript writing, review or editing, and approved the final submitted version of the manuscript.

## Data availability

Additional data and supportive evidence are available in Supplementary Information. Other requests can be directed to the corresponding author.

## Competing interests

The authors declare no competing interests. Funders had no role in design, interpretation and analysis of the presented results, decision to publish or manuscript preparation.

## Notes

### Competing Interest Statement

The authors have declared no competing interest.

