## Supplementary Information for "N-formylkynurenine but not kynurenine enters a nucleophile-scavenging branch of the immune-regulatory kynurenine pathway"

#### 1. Supplementary methods

**Metabolite identification on liquid chromatography - high-resolution mass spectrometry (HPLC-HR-ESI-MS)**

To identify molecular masses of metabolites with high accuracy, an Agilent 1100 HPLC/6530B Q-TOF ESI MS system was used (results show at Supplementary Fig. 3). The mobile phases consisted of A = 10 mM ammonium formate (pH 4) and B = 80% acetonitrile supplemented with 10 mM ammonium formate (pH 4). They were mixed as follows: 0 - 2 min (94.3% A), 6.5 – 7.5 min (19.2% A), 8 – 10.5 min (94.3% A). The MS acquisition conditions were: 50-2000 m/z range, 0.99 spectra/second, 100 V fragmentor voltage, 65V skimmer voltage. Ionisation conditions were: 275°C drying gas temperature, 10 L/min gas flow, 40 psi nebulizer pressure, 3500 V positive/negative capillary voltage, 500 V positive/negative nozzle voltage.

### 2. Supplementary figures

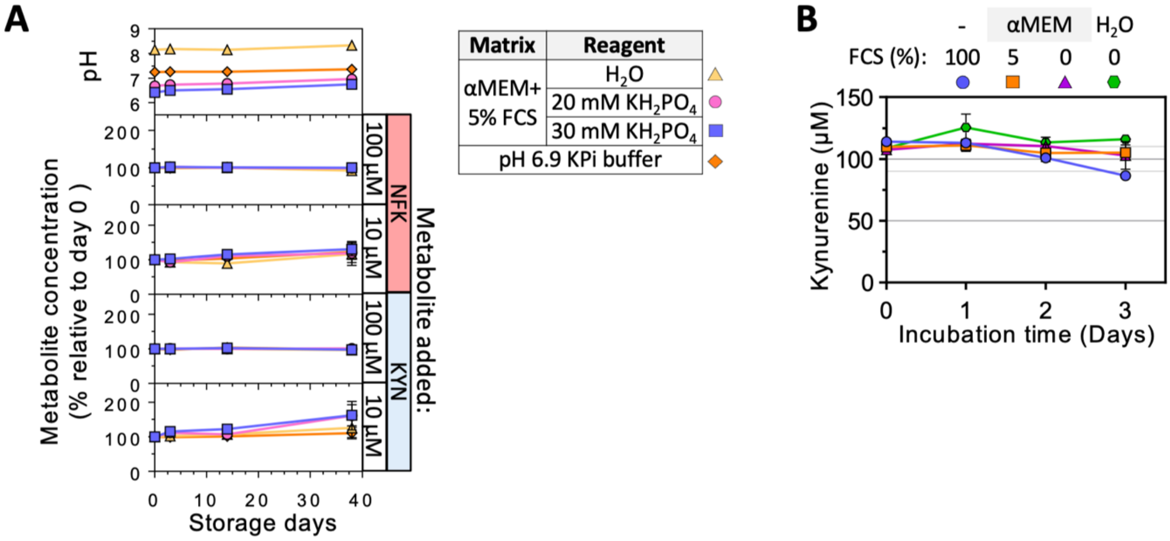


**Supplementary Fig. 1.**

**(A) Stability of NFK and KYN at two concentrations (rows) at -80°C storage in 3 different environments (symbols and colours).** Assembled samples (10 μL metabolite into 110 μL KPi buffer or pre-incubated αMEM matrix) were aliquoted (50 μL) and neutralised using reagents (right-side annotation table). Neutralised samples were stored at -80°C, and at each time point, samples were thaw, measured pH with an electrode, and analysed by HPLC-UV. Symbols connected by solid lines represent mean concentration or pH values normalised to Day 0 values from 2 independent experiments. Whiskers are SEM. Annotation panels on the right indicate metabolites and concentrations added into matrices.

**(B) Stability of KYN at 37°C in 4 different pH 7.4 bicarbonate buffered environments (symbols and colours).** Points connected by solid lines and whiskers are the mean and SEM of KYN concentration from two experiments with technical replicates, respectively.

<u>Abbreviations</u>: NFK: N-formylkynurenine; KYN: kynurenine; FCS: foetal calf serum; αMEM: alpha Minimum Essential Medium; KPi: potassium phosphate buffer; $KH_2PO_4$: potassium dihydrogen phosphate.

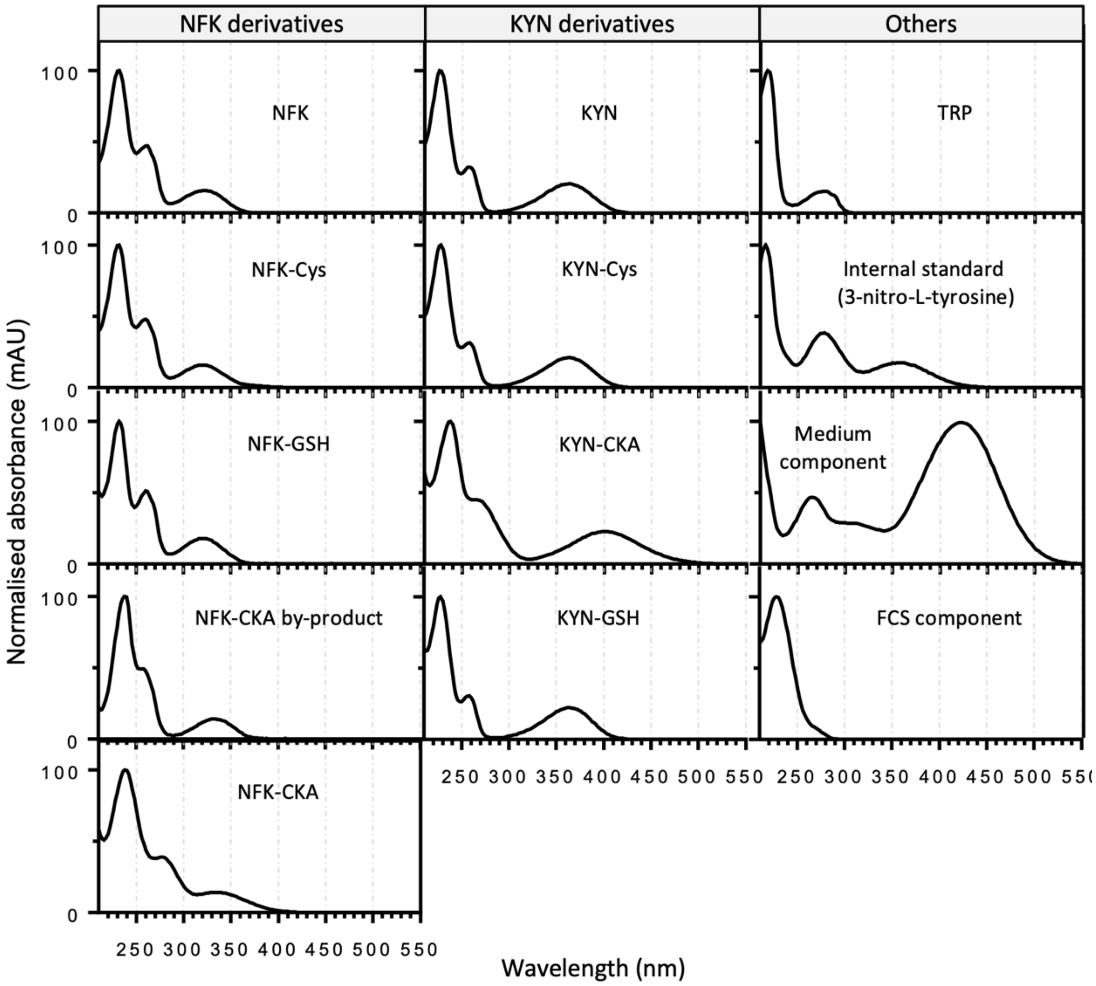


**Supplementary Fig. 2. Normalised absorbance spectra of principal peaks (>10% of the NFK peak height) identified from the chromatogram in Fig. 3 and 4**. Absorbance spectrum of each principal peaks at their retention time from HPLC-DAD was normalised to the highest and lowest absorbance signal detected within the range from 210 to 550 nm.

Abbreviations: TRP: tryptophan; Cys; cysteine; GSH: glutathione; CKA: carboxyketoalkene.

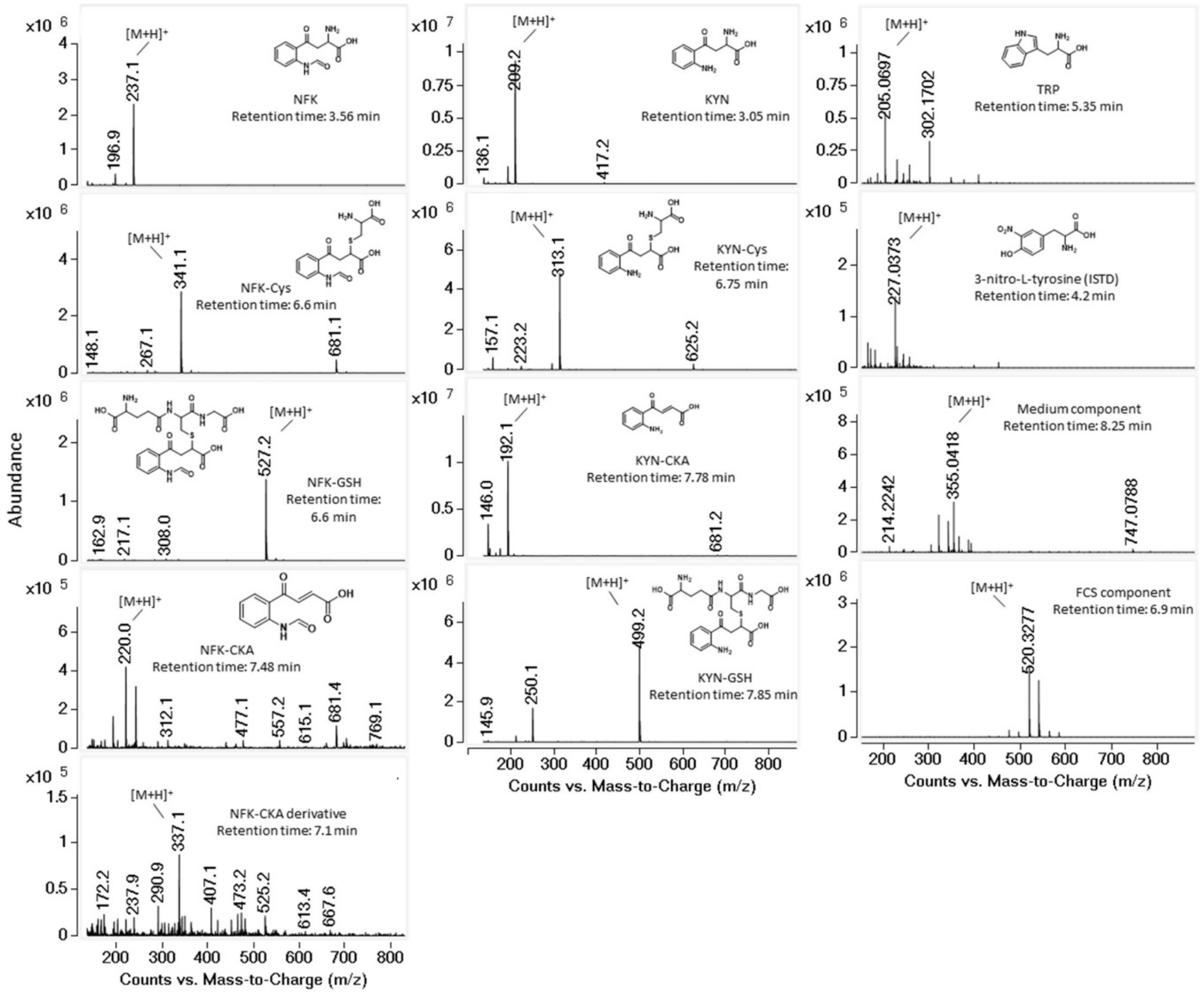


**Supplementary 3. Mass spectra (135 to 800 *m/z* range) of principal peaks (>10% of the NFK peak height)** at their retention time by HPLC-ESI-MS. The protonated molecule $[M+H]^+$ was labelled, with the retention time from HPLC-DAD and chemical structure if known. The mass spectrum of NFK and KYN derivatives were generated from HPLC-ESI-MS (section 4.3.3) and others were from HPLC-HR-ESI-MS (section 5.1).
